# Intercellular calcium signaling is regulated by morphogens during *Drosophila* wing development

**DOI:** 10.1101/104745

**Authors:** Pavel A. Brodskiy, Qinfeng Wu, Francisco J. Huizar, Dharsan K. Soundarrajan, Cody Narciso, Megan K. Levis, Ninfamaria Arredondo-Walsh, Jianxu Chen, Peixian Liang, Danny Z. Chen, Jeremiah J. Zartman

**Author notes:** These authors contributed equally to this work (Pavel A. Brodskiy, Qinfeng Wu). Email for correspondence Phone number for correspondence.

## Abstract

Organ development is driven by a set of patterned inductive signals. However, how these signals are integrated to coordinate tissue patterning is still poorly understood. Calcium ions (Ca^2+^) are critical signaling components involved in signal integration and are regulated by a core Ca^2+^ signaling toolkit. Ca^2+^ signaling encodes a significant fraction of information in cells through both amplitude and frequency-dependent regulation of transcription factors and key regulatory enzymes. A range of intercellular Ca^2+^ transients, including coordinated oscillations, recently have been reported in *Drosophila* wing discs. In an accompanying paper, we show that impaired Ca^2+^ signaling impacts the final size and shape of the wing. Here, we discover specific spatiotemporal signatures of Ca^2+^ transients during wing disc development. To do so, we developed a new neural-network-based approach for registration of oscillatory signals in organs that frequently move during imaging, and a pipeline for spatiotemporal analysis of intercellular Ca^2+^ oscillations. As a specific test case, we further demonstrated that the morphogen pathway, Hedgehog, controls frequencies of Ca^2+^ oscillations uniformly in the tissue and is required for spatial patterning of oscillation amplitudes. Thus, the time-averaged dynamics of spontaneous intercellular Ca^2+^ transients reflect the morphogenetic signaling state of the tissue during development. This suggests a general mechanism of physiological signaling that provides a memory of morphogenetic patterns. Additionally, our study provides a powerful approach for registering and quantifying oscillatory dynamics in developing organs.

## Introduction

The mechanical and biochemical processes critical for organogenesis require coordination of cell activities. This coordination is facilitated by morphogens, which are diffusible factors whose concentration conveys positional information to cells. The *Drosophila* wing imaginal disc pouch has been used to study how a simple sheet of epithelial cells is sculpted into the intricate structure of an adult wing (Bier, 2005; Blair, 2007; Hariharan, 2015; Neto-Silva et al., 2009; Restrepo et al., 2014; Singh and Irvine, 2012) (Fig. 1A-B). In the larval state, morphogens divide the wing disc pouch into regions that define the differentiation state of cells and coordinate morphogenesis (Fig. 1A). Hedgehog (Hh) and Decapentaplegic (Dpp) signaling patterns the anterior (A)/posterior (P) axis, while Wingless (Wg) signaling patterns the dorsal (D)/ventral (V) axis. However, the logic of morphogen signal integration and memory is still largely unclear.

**Figure 1:**
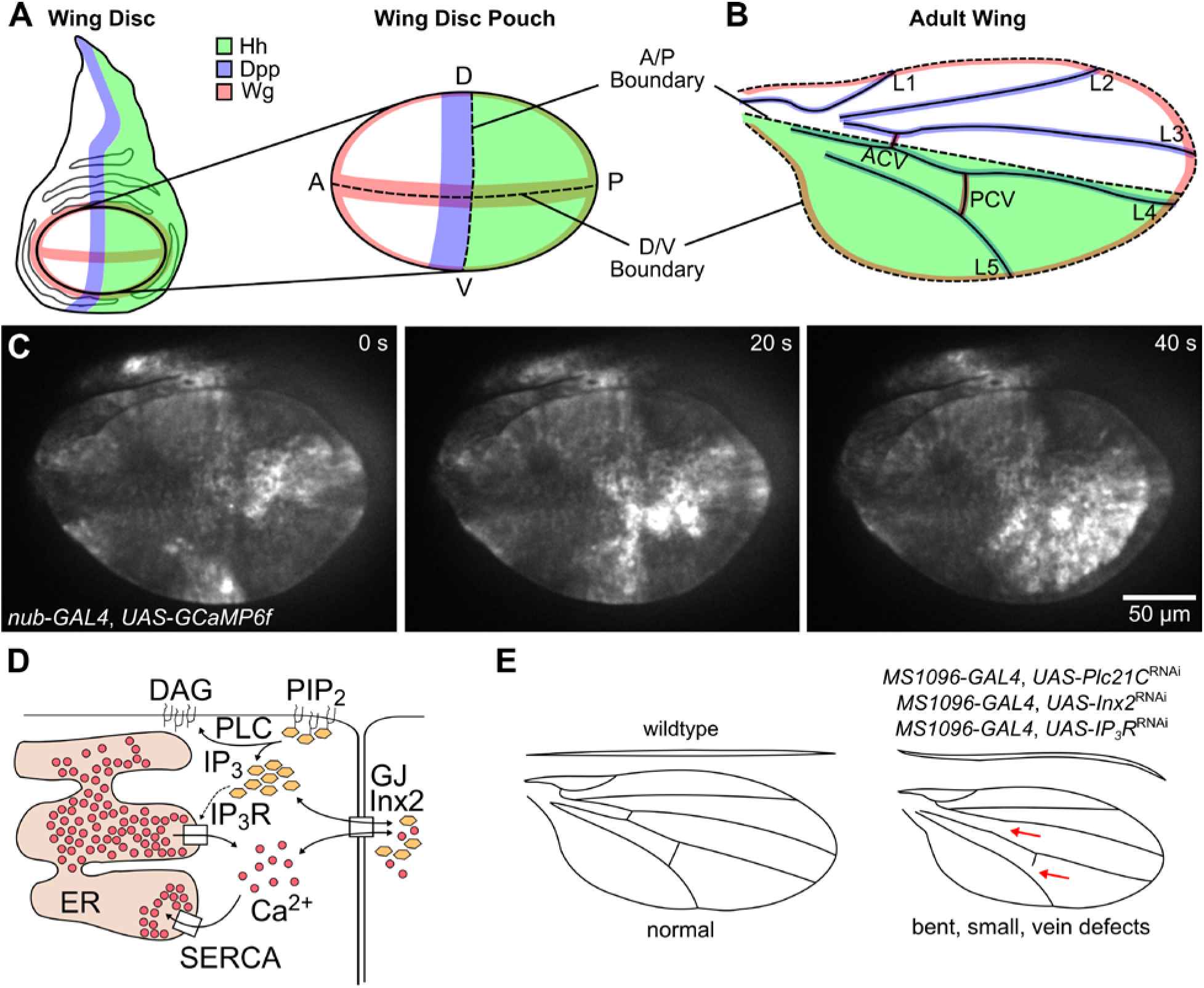
The wing imaginal disc is a model system of organ growth and patterning and is characterized by intercellular Ca^2+^ transients that propagate throughout the tissue. (A-B) Illustration of (A) the larval wing disc pouch, which is a patterned monolayer epithelium that will form (B) the adult wing. Spatial patterning of cell fates is determined by the combined actions of locally secreted morphogenetic signals such as Hedgehog (Hh), BMP/Decapentaplegic (Dpp), and WNT/Wingless (Wg). The central ellipsoidal region of the wing disc is the pouch and is subdivided into four quadrants defined by intersecting dorsal – ventral (D/V) and anterior-posterior (A/P) axes. Hh specifies the anterior and posterior (green) compartments and defines location of Dpp generation and secretion. The source of Wingless in the pouch is along the dorsal – ventral boundary, which defines the wing margin. (C) *Ex vivo* confocal imaging of relative concentrations of cytosolic Ca^2+^ with the genetically encoded Ca^2+^ sensor GCaMP6 (Tian et al., 2009, 6), expressed using the Gal4/UAS system (*genotype: nub>GCaMP6f*). Live imaging of explant cultures enables high resolution imaging of Ca^2+^ signaling for extended periods (multiple hours). (D) Mechanism for Ca^2+^ signaling in the wing disc. Phospholipase C (Plc21C) generates 1,4,5-trisphosphate (IP_3_), which triggers the release of Ca^2+^ from the endoplasmic reticulum (ER). (E) Illustration of adult wing with perturbations to IP_3_R signaling. Inhibition of the components of Ca^2+^ signaling results in small, bent wings with cross vein defects as we show in our accompanying paper (DEVELOP 2017/159251).

Recently, intercellular Ca^2+^ transients (ICTs) have been observed in the wing disc in vivo and ex vivo (Fig. 1C). They have been implicated in ensuring robustness in regeneration (Restrepo and Basler, 2016), tissue homeostasis (Balaji et al., 2017), and mechanotransduction (Narciso et al., 2017). Ca^2+^ signaling in the wing disc relies on a highly-conserved Ca^2+^ signaling toolkit. (Fig. 1D). Ca^2+^ signaling encodes information through the Ca^2+^ signature, which includes the amplitude (Amp), frequency (F), width at half max (WHM), and basal Ca^2+^ concentration level (B) (McAinsh and Pittman, 2009). In an accompanying paper, we show that inhibiting key components of the intercellular Ca^2+^ toolkit leads to smaller, bent wings with missing cross veins (DEVELOP 2017/159251, Fig. 1E). Cross vein specification requires precise regulation by morphogens (Fig. 1B). However, it is unknown whether Ca^2+^ signaling is a random phenomenon or spatially patterned in the wing disc.

Here we develop a novel pipeline for registration and spatiotemporal analysis of tissue-scale Ca^2+^ signaling. We test this pipeline on a large set of ex vivo imaging data to identify several signatures of Ca^2+^ signaling that are regulated by morphogens. Spatial patterning of the amplitude of oscillations requires Hh signaling, and Hh signaling suppresses the frequency of Ca^2+^ oscillations in the wing disc pouch. This study further establishes a model whereby morphogenetic signaling impacts the physiological network that regulates Ca^2+^ signaling. We posit that this secondary level of patterning in turn is important for robust tissue morphogenesis and regeneration.

## Results and discussion

### A novel pipeline for spatial analysis of Ca^2+^ oscillations

We used a *nub-GAL4, UAS-GCaMP6* reporter line to visualize Ca^2+^ in the wing disc pouch (Narciso et al., 2017). *GCaMP6* is a fluorescent sensor for Ca^2+^, and the *nub* driver is uniformly expressed in the wing disc pouch. Quantitative spatial analysis of in vivo Ca^2+^ is challenging because of larval motion and optical interference from the cuticle. In vivo observations of oscillations are limited to short term experiments due to the need to keep photo-sensitive larvae constrained without significantly impacting cellular processes. These challenges led us to develop an ex vivo imaging and analysis pipeline (Fig. 2). We used ex vivo culture in the chemically-defined ZB media to reduce the effect of batch variability (Burnette et al., 2014). Although the accompanying paper shows that ZB media with 2.5% fly extract best recapitulates in vivo Ca^2+^ dynamics (DEVELOP 2017/159251), we found that 15% fly extract was ideal to yield oscillatory signals of sufficient strength for reliable quantification.

**Figure 2:**
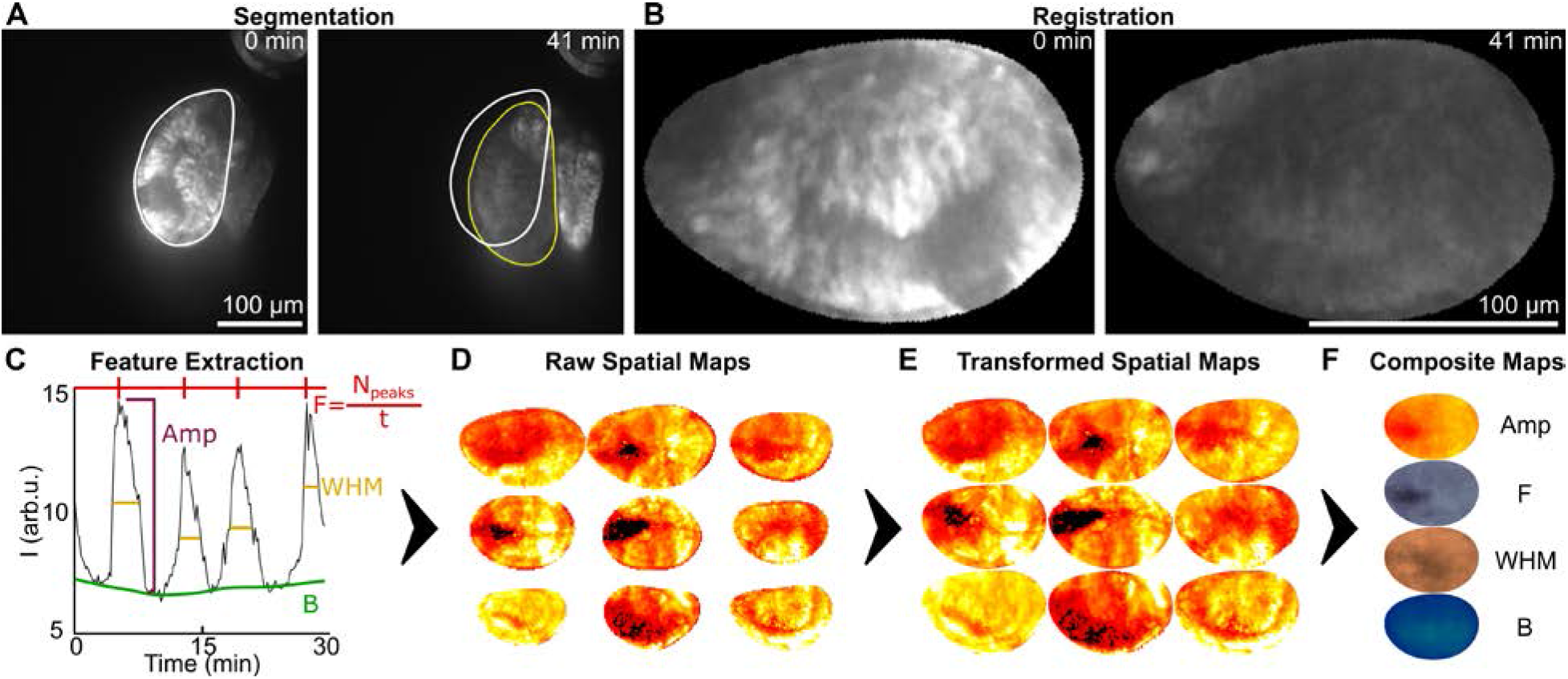
Pipeline for spatiotemporal characterization of Ca^2+^ signatures. (A) Automatic segmentation of pouch region with deep-learning segmentation algorithm. Outlines represent the pouch at the initial time-point (white) and the final time-point (yellow). (B) Registration of raw images onto canonical pouch shapes. (C) Extraction of amplitudes (Amp), frequency (F), width at half max (WHM) and basal level (B) from Ca^2+^ trace for individual regions-of-interest (ROI). (D) The Ca^2+^ signature is computed for each spatial position within each pouch to generate a spatial map. (E) Each spatial map is transformed onto a canonical coordinate system to align the A-P and D-V axes. (F) Each set of spatial maps is averaged at each position to generate a composite spatial map.

A common roadblock for whole-organ analysis of fast signaling dynamics is tissue registration across frames within a movie and across multiple samples of different shapes and sizes. The pouch region of the wing imaginal disc develop into the adult wing. However, no GAL4 driver exists that is uniformly expressed throughout only the wing disc pouch. We used the nubbin GAL4 driver, which is also expressed in a small region of the hinge (Fig. 2A). We developed a segmentation pipeline to segment the pouch from a single channel provided by the GCaMP6 signal. This pipeline detects the borders of the pouch, and transforms the underlying images so that the edges of the pouch are the same from frame to frame. Segmentation was done using a deep neural network approach, with training data from 800 images of wing disc collected under a variety of genetic and culture conditions (Fig. 2A). B-spline based registration was used to transform images onto a shape defined by the first time-point (Fig. 2B). The pipeline ensures coordinates within the pouch are consistent from frame to frame, allowing spatial analysis of Ca^2+^ signatures.

We divided the wing disc pouch into square regions of interest (ROIs) of 2.8 μm. We used average intensity within each ROI over time to obtain Ca^2+^ signaling signatures (Fig. 2C). Signatures were obtained for each ROI and stored as a spatial map (Fig. 2D). These spatial maps were then transformed such that the A/P and D/V axes were consistent across samples within a category (Fig. 2E). Finally, each set of spatial maps was averaged to obtain a composite spatial map for each feature (Fig. 2F). A fraction of the wing discs exhibited a “fluttering” phenotype as described in our accompanying paper (DEVELOP 2017/159251). In fluttering discs, Ca^2+^ signaling is so over-stimulated that cells stay at high levels of Ca^2+^, with higher-frequency, lower-amplitude oscillations than non-fluttering tissues. Fluttering videos were omitted from quantitative analysis because they do not provide sufficient information on basal intensity.

### Spatial patterning of Ca^2+^ signatures emerges throughout development

Surprisingly, spatial analysis of Ca^2+^ signaling revealed a difference in amplitude between the anterior (A) and posterior (P) compartments of the pouch (Fig. 3A), suggesting spatial patterning of Ca^2+^ signatures. Across development, 91% of discs had higher average amplitudes (Amp) in the P compartment, which is where Hh (Fig. 1A) and Engrailed are expressed. The P compartment, however, is not sensitive to Hh signaling (Robbins et al., 2012). The average Amp is approximately 33% higher in the P compartment than the A compartment (Fig. 3B). The basal level was lower along the A-P axis, and higher along the D-V axis. Of note, the increase in basal concentrations of Ca^2+^ along the D-V axis correlates with tissue thickness (Widmann and Dahmann, 2009). The slight decrease in basal level along the A-P axis correlates with a slight decrease in height that can become a fold when Dpp signaling is suppressed (Shen et al., 2008; Umemori et al., 2007). In discs that were not fluttering, we did not observe any spatial difference in frequency. However, for fluttering discs we qualitatively observed that the anterior half of the D-V axis often exhibited lower-frequency, higher-amplitude oscillations unlike the rest of the tissue as described in our accompanying paper (DEVELOP 2017/159251). Taken together, these findings show that these features of Ca^2+^ signaling are highly-correlated with three spatial patterns of the tissue that are downstream of the morphogen signaling pathways: Hh, Dpp, and Wg signaling.

**Figure 3:**
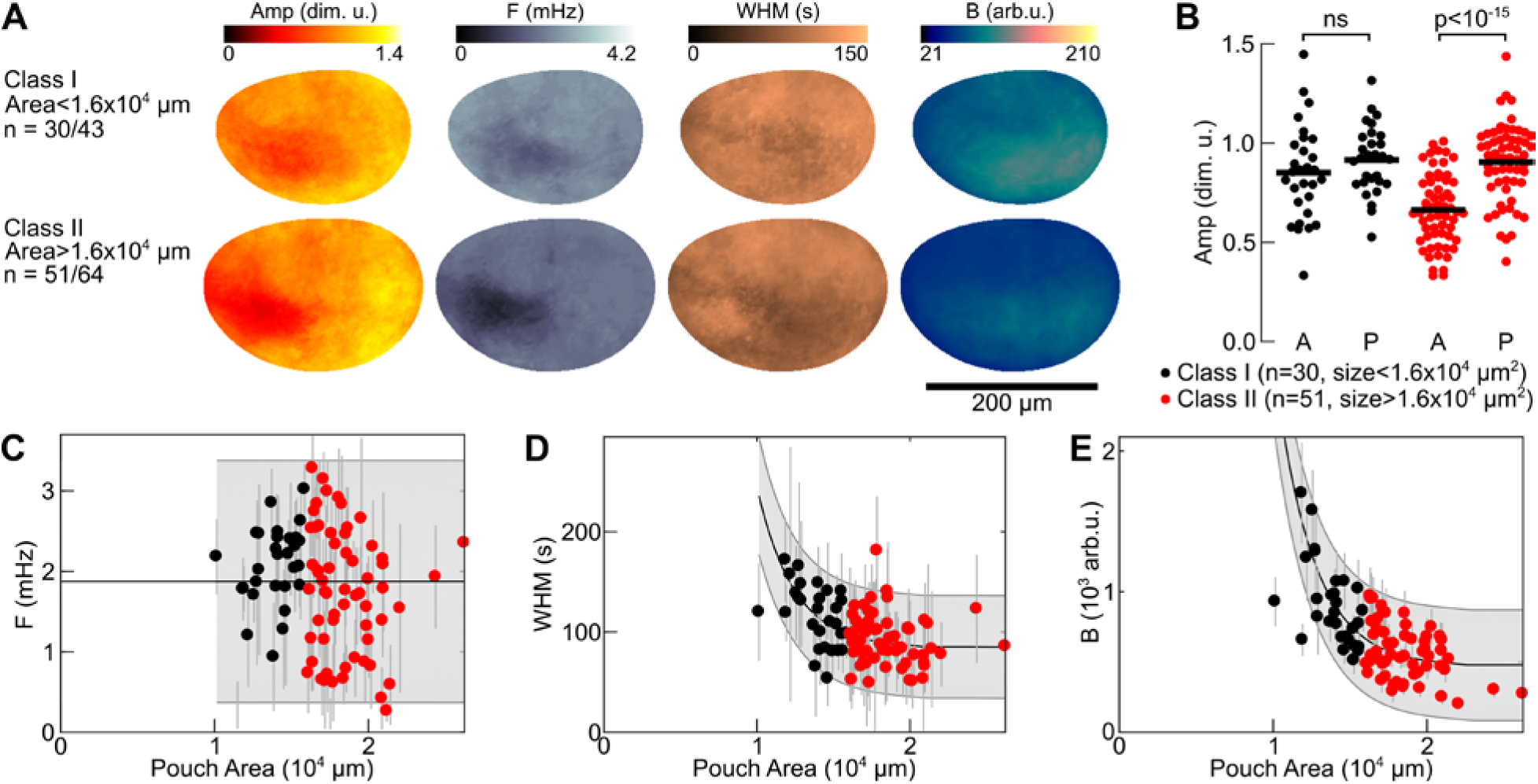
ICWs are spatiotemporally patterned during development. (A) Composite spatial maps of the median ICW summary statistics. Pouches were grouped by pouch size for comparison of smaller “younger” discs to “older” larger discs. Ca^2+^ signaling in *nub-GAL4; UAS-GCaMP6f* pouches followed one of two patterns: small pouches with higher WHM and basal intensity, and larger pouches with a higher Amp in the P compartment. (B) Average amplitude (Amp, dim. u.) for each compartment was greater in the P compartment for the class of larger pouch. (C) Frequency, F (mHz) did not change over development. (D) Width at half max, WHM (s) decreased slightly over development (for exponential fit, adjusted R^2^=0.15, p<3x10^−3^ by F-test). (E) Basal intensity, B (arb. u.) decreased over development (for exponential fit, adjusted R^2^=0.77, p<1x10^−17^ by F-test). Error bars represent standard deviations within each spatial map. Grey shaded area is the 95% confidence interval of prediction. p values were obtained by paired t-test.

We examined Ca^2+^ signatures in pouches of different developmental ages to investigate developmental patterning of Ca^2+^ signatures. We grouped the wing discs by pouch size, as a proxy for developmental stage of the organ (Bittig et al., 2009; Hamaratoglu et al., 2011; Hufnagel et al., 2007; Wartlick et al., 2011) (Fig. 3). We found that the difference in amplitudes between compartments increases with increasing pouch size (Fig. 3A, B). The frequency was independent of pouch size (for linear or exponential fit, adjusted R^2^<0.05, Fig. 3A, C). The width at half max is weakly correlated with pouch size, and was not spatially patterned between compartments in the pouch (Fig. 3A, D). Basal concentration of Ca^2+^ decreases with increasing pouch size (Fig. 3A, E). Averaged over the compartment level, frequencies are only slightly different (Fig. 3A). We suggest that frequency may be driven primarily by diffusion within the whole tissue, while amplitude is driven by the difference between Ca^2+^ concentration in the cytoplasm and ER of individual cells (Fig. 3A, C).

Differential Ca^2+^ oscillations could be caused by difference in growth dynamics and metabolic activities between the two compartments (Martín and Morata, 2006). In our accompanying paper, we show that frequency decreases with age in staged larvae from 5 to 8 days after egg laying (DEVELOP 2017/159251). Combined with our finding that smaller pouches exhibit more fluttering (30% vs 20%), and have higher basal levels of Ca^2+^, this suggests that Ca^2+^ signaling decreases with developmental age. ICTs could then be a phenotypic response to growth and represent stress dissipation that occurs as the tissue size increases as recently proposed (Narciso et al., 2017). These results show that the overall basal level decreases as the discs mature and that a spatial patterning of ICT amplitudes across the posterior half emerges at later stages of development.

### Suppression of Hedgehog signaling increases Ca^2+^ activity

The discovery of A/P patterning of average ICT amplitudes suggest that morphogen pathways shape Ca^2+^ dynamics. As a test case, we perturbed the Hh signaling pathway, which directs the development of the A-P compartment boundary and is upstream of Dpp signaling (Akiyama and Gibson, 2015; Basler et al., 1994; Dahmann et al., 2011; Lecuit et al., 1996; Rodriguez and Basler, 1997). Hh signaling activity is dependent on the protein Smoothened (Smo), whose activity is inhibited by the receptor Patched (Ptc) (Robbins et al., 2012). In the absence of Hh, the full-length transcriptional factor Ci (Ci^155^) is truncated into a repressor form, Ci^R^ (Robbins et al., 2012). Binding of Hh to Ptc blocks Ptc-mediated Smo inhibition, allowing Smo to stop Ci cleavage (Robbins et al., 2012). We can activate the pathway through expression of *ci(m1-3)*, a constitutively-active form of Ci (Chen et al., 2000). *ci(m1-3)* has three mutated PKA phosphorylation sites in the activation domain, so it cannot be truncated.

We analyzed ICWs with upregulated and downregulated Hh signaling to determine the effect of perturbations on ICWs. We used *nub-GAL4, UAS-GCaMP6f/UAS-ci(m1-3)* to activate the Hh pathway, and *nub-GAL4, UAS-GCaMP6f/UAS-smo^RNAi^* to suppress the Hh pathway in the wing disc pouch. The perturbations were validated by examination of the wing phenotypes (Table S1). We found that suppression of Hh signaling was sufficient to abolish spatial patterning of ICW amplitudes between the A and P compartments and leads to a higher frequency in the entire pouch (Fig. 4B). Activation of Hh signaling resulted in a lower frequency in the entire disc than *nub-GAL4, UAS-GCaMP6f* (Fig. 4B). These results demonstrate that Hh signaling negatively regulates Ca^2+^ oscillation frequencies in the wing disc pouch, and that spatial patterning of the amplitude of Ca^2+^ oscillations is downstream of Hh signaling.

**Figure 4:**
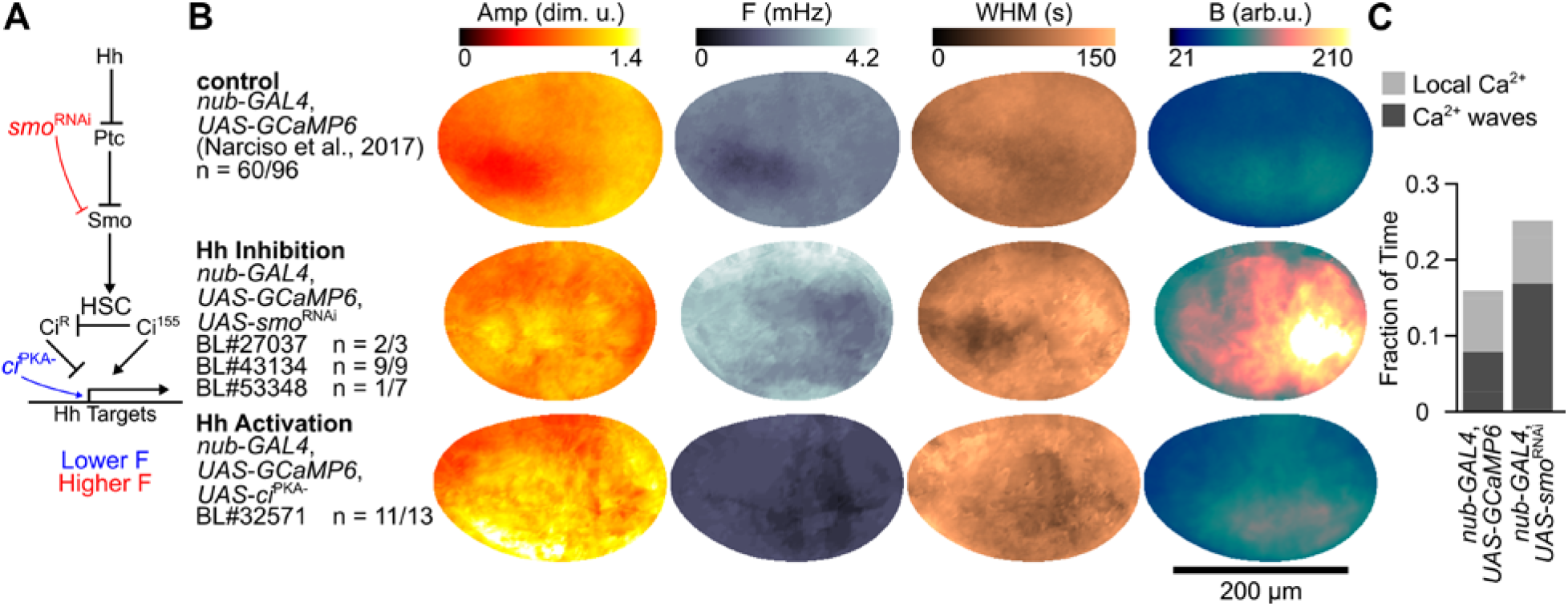
Hedgehog signaling suppresses frequency of Ca^2+^ oscillations and patterns amplitude. (A) Flowchart of the Hh signaling pathway. Hh binds to and inhibits patched (Ptc), which causes sequestration of the GPCR smoothened (Smo). When Smo is not sequestered, it prevents the truncation of Ci into its repressor form, Ci^R^. Ci^PKA-^ is a constitutively active form of Ci. Ci is a transcription factor for a group of genes and the untruncated form Ci^155^ induces transcription, whereas the truncated form Ci^R^ suppresses transcription. (B) Composite spatial maps of the median ICW summary statistics for the tester line crossed with either itself, a Hh inhibitor, or a Hh signaling activator. Sample size represents number of analyzed videos over total number of videos, counting fluttering discs. Inhibition of Hh signaling led to an increase in frequency (F) and basal level (B), and activation of Hh signaling led to a decrease in F. Both perturbations led to a loss of A-P patterning of amplitude (Amp). (C) Quantification of Ca^2+^ activity in vivo. The fraction of time that discs are undergoing each type of signaling is reported. Local Ca^2+^ events include spikes of Ca^2+^ within a cell or small transients travelling over fewer than five cell diameters. Ca^2+^ waves indicate longer-range Ca^2+^ oscillations including fluttering due to overstimulation. Inhibition of Hh signaling doubles the frequency of long-range Ca^2+^ oscillations in the wing disc in vivo.

Further, we performed in vivo imaging of *nub-GAL4, UAS-GCaMP6f* crosses to qualitatively test the effect of Hh signaling perturbations on Ca^2+^ during development. We quantified the ratio of wing discs undergoing either local Ca^2+^ events, Ca^2+^ waves, or no activity. We found that discs expressing *UAS-Smo^RNAi^* exhibited long-range waves 17% of the time (n=40) compared with 8% of the time for the tester line (Fig 4C, n=102). Local Ca^2+^ events such as spikes and small transients were roughly the same between the two conditions (8.3% and 8.0% respectively). These findings show that Hh signaling also modulates Ca^2+^ signaling in vivo by both increasing the duration and extent of Ca^2+^ signaling events.

One possible explanation for the patterning of Ca^2+^ oscillations is that Hh signaling regulates patterning through actomyosin-mediated mechanical stresses. Hh pathway regulates tissue bending, folding, and invagination through actomyosin in *Drosophila* eye discs (Corrigall et al., 2007), embryo (Czerniak et al., 2016) and fore-gut (Lechner et al., 2007). We have shown that ICWs can be stimulated by the release of compression in wing discs (Narciso et al., 2017). Laser ablation of cells leads to rapid dissipation of mechanical stress and Ca^2+^ flashes in the wing discs cultured in fly extract-free medium (Narciso et al., 2015). In addition, it was shown that the myosin II inhibitor blebbistatin interrupted Ca^2+^ oscillations in the wing disc (Balaji et al., 2017). High levels of Hh signaling have been shown to induce folds, which can be prevented by expression of Dpp (Shen et al., 2008; Umemori et al., 2007). Taken together, Hh signaling may regulate ICT patterning by differential regulation of actomyosin in different compartments. Perturbations to morphogen signaling lead to dysregulation of Ca^2+^ signaling, with severe implications for morphogenesis. These effects are analyzed in further detail in our accompanying paper (DEVELOP 2017/159251) and have many implications for understanding the progression of birth defects and diseases.

### Conclusions

Here, we report a novel approach for exploring relationships between Ca^2+^ signaling and morphogen signaling. Through this approach we have used this pipeline to show that time-averaged Ca^2+^ signaling dynamics are spatially patterned. Spatial composite maps of ICW features reveal A/P patterning of average ICT amplitudes and temporally decreasing basal intensity as wing discs reach their terminal size before pupariation. Thus, time-varying Ca^2+^ signals in cells are capable of multiplexing a wide range of morphogenetic signals during organ development. These results suggest a model where each morphogen signaling pathway impacts Ca^2+^ signaling signatures uniquely.

## Materials and Methods

### Fly strains and genetics

A *nub-GAL4, UAS-GCaMP6f* reporter line was used to measure relative Ca^2+^ signals in the wing disc pouch. Gene perturbations to the core Ca^2+^ and Hedgehog signaling pathways (UAS-*Gene X*^RNAi^ or UAS-*Gene X* (overexpression) were crossed to a genetic tester line (*nub-GAL4, UAS-GCaMP6f/CyO*) that enables combined visualization of Ca^2+^ signaling with down regulation of genes through expression of RNAi in the wing disc. When possible, multiple independent RNAi lines were tested for each gene investigated (Table S1, 4). Stocks are obtained from Bloomington *Drosophila* Stock Center as indicated by stock number (BL#). Progeny wing phenotypes are from F1 male progeny emerging from the nub-Gal4, UAS-GCaMP6f/CyO x UAS-X cross and summarized in Table S1. The tester line (w1118; nubbin-GAL4, UAS-GCaMP6f/CyO) was generated in (Narciso et al., 2017). Flies were raised at 25°C and 12-hour light cycle.

### In vivo imaging setup

Wandering 3^rd^ instar *nub-GAL4, UAS-GCaMP6f/CyO* larvae were collected for imaging and rinsed vigorously in deionized water. They were dried completely and then adhered to a coverslip for imaging with scotch tape covering the larvae. The larvae were attached with their spiracles facing toward the coverslip to align the wing discs toward the microscope. An EVOS Auto microscope was used to image the larvae. The larvae were imaged at 20x magnification for 20 minutes. Images were taken every 15 seconds.

### Organ culture media

ZB media + 15% fly extract contains 79.4% (v/v) ZB media, 0.6% insulin, 15% ZB-based fly extract and 5% penicillin/streptomycin. ZB media was developed as a chemically defined media to support *Drosophila* cell culture (Burnette et al., 2014), and was used as the basal media for all experiments. ZB-based fly extract is undefined serum extracted from ground flies using ZB media as the base. It is commonly used in culture media as a substitute of fetal bovine serum (FBS) to support *Drosophila* tissue growth and prolong the culture time of cultured wing discs. ZB-based fly extract is made from the following protocol: One gram well-nourished mature flies, were homogenized in a tissue homogenizer (15 ml capacity, 0-1 mm clearance) with 6.82*x* ml of ZB media. This homogenate was centrifuged at 2600 rpm for 20 min at 4°C. The supernatant and the oily film above it were removed and heat-treated at 60°C for 20 min. This preparation was then spun at 2600 rpm for 90 min at 4°C. The supernatant (fly extract) was removed, sterilized by 0.2 μm filtration, and stored at 4°C.

### Wing disc imaging setup

Wing discs were dissected from 3^rd^ instar larvae and cultured and imaged in the organ culture media on a cover slip based on our previously developed protocol (Zartman et al., 2013). A cell culture insert (EDM Millipore), after truncated the legs, was put on top of the pool of media to immobilize the wing discs. 50 μL of embryo oil was added along the outer periphery of the insert to seal. 100 μL of *organ culture* media was added on top of membrane of the insert. The setup was then transferred to a confocal microscope for Ca^2+^ activity imaging. Imaging was performed on a Nikon Eclipse T*i* confocal microscope (Nikon Instruments, Melville, NY) with a Yokogawa spinning disc and MicroPoint laser ablation system (Andor Technology, South Windsor, CT). Image data were collected on an iXonEM+ cooled CCD camera (Andor Technology, South Windsor, CT) using MetaMorph^®^ v7.7.9 software (Molecular Devices, Sunnyvale, CA). All experiments were performed immediately following dissection to minimize time in culture, except for drug experiments that were incubated for one hour in drug solution or vehicle carrier control prior to imaging. Discs were imaged at a three z-planes with a step size of 10 μm, 20x magnification and 10-second intervals for a total period of one hour, with 200 ms exposure time, and 50 mW laser, and 44% to 70% laser intensity. We found that the GCaMP sensor saturated at roughly 80% laser intensity, but that intensity was linearly related at values under 70% (Fig. S5). Image intensity was linearly normalized to be comparable at 50% laser intensity.

### Data pre-processing

Microscopy resulted in 4D time-lapse data (512 pixels by 512 pixels by 3 z-planes by 361 time points). The z-stack data was max-projected in FIJI (Schindelin et al., 2012) to yield z-projected time-lapse videos. Discs that moved over the imaging session by more than several cell diameters were registered with the *StackReg* function in FIJI. Time points were selected such that discs were only analyzed during times in which discs were immobile, with the shortest time-lapse analyzed being 20 minutes.

### Image segmentation

For the image frame at each time point, an ROI defining the pouch was obtained using a novel deep-learning based segmentation algorithm that we developed. Each frame was segmented with a fully-convolutional neural network (FCN) module, and the ROI boundaries were refined by a graph-search algorithm. The FCN module was trained on 800 images of wing discs expressing *nub-GCaMP6* with various stages of Ca^2+^ activity. The FCN was used to segment only the pouch region, and not the *nub*-expressing region dorsal to the pouch. Note that traditional based segmentations that we experimented with could not distinguish between these two regions well enough.

### Image registration

The first ROI in an image video was taken as the reference ROI and was rotated such that the major axis was horizontal. Each image frame was transformed using a novel registration algorithm that we developed such that all pouch images had the same final dimensions. Note that commonly-used Intensity-based mapping algorithms are not suitable for registration of tissues with Ca^2+^ waves, as the highly-transient Ca^2+^ signaling activity is often a stronger landmark than the underlying tissue. Therefore, we used a custom method for transforming wing disc pouches onto a shape based solely on the ROI produced by segmentation. First, a rigid transformation was applied to each image to roughly align the ROIs of the images in the sequence with the reference ROI. Next, B-spline free-form deformation (FFD) was used to transform each frame such that the ROI for the frame matched the reference ROI.

### Identification of the axes

The posterior (P) and dorsal (D) compartments were identified manually based on the characteristic shapes of pouch (Fig. 1A, Fig. S3A). Pouches were flipped such that the A compartment was on the left, and the D compartment was on the top. A custom MATLAB GUI was used to reduce error in manual pouch orientation (Fig. S3A). Each image was presented as a 2x2 grid of the image either not transformed, flipped left/right, flipped top/down, or rotated 180 degrees in random order. The user selected the correctly-oriented pouch. The order in which the pouches were displayed was randomized, and a consensus was reached once three guesses were made if all three were the same. 77% of the samples were unambiguously classified in the first attempt in this way. If the first three guesses were not the same, then when more than half of the guesses were the same orientation, the most common orientation was taken to be the consensus. Two images were removed from the analysis because no consensus was reached after seven attempts. For each pouch, the A-P and D-V axes were identified manually by segmentation with a custom MATLAB GUI (Fig. S3B).

### Feature extraction

For each pouch sample, signals were obtained by taking average intensity (F(t)) from 4x4 pixel (2.8 μm x 2.8 μm) spatial bins arrayed with square packing across the segmented disc pouch (Fig. S1A-C). Video durations ranged from 20 to 60 minutes. Each signal was decomposed into the following features: amplitude, frequency, width at half max, and basal intensity, which comprise the Ca^2+^ signature of the ROI. Amp is defined as the mean of the amplitudes of the peaks, where the amplitude of each peak is the prominence of the peak in the *findpeaks* algorithm. F (mHz) is the number of peaks detected divided by the length of the signal in time. WHM (s) is defined as the time that a peak is above the peak value minus half the prominence. B (arbitrary units) represents the equilibrium cytoplasmic concentration of Ca^2+^, and is the minimum of the signal (Fig. 2, Fig. S1D-E).

The normalized intensity (ΔF(t)/F_o_) was approximated by using a bandpass Gaussian filter, where the larger sigma value adjusted for change in basal level over time, and the smaller sigma value compensated for stochastic noise. Spikes in signaling activity were extracted from ΔF(t)/F_0_ using the MATLAB *findpeaks* algorithm, with a minimum amplitude (Amp_min_), refractory period, and width at half max (WHM_min_), for a total of five parameters. MATLAB’s genetic algorithm *ga* was used to calibrate the feature extraction parameters (Fig. 2, Fig. S2, Table S2). To generate reference values for optimization of feature extraction parameters, 233 signals (I(t)) were randomly selected from 656,000 total signals, and manually annotated to identify the times (t_1_ and t_2_) at which each peak begins and ends. From this, the basal level was taken to be:

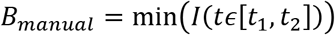

the amplitude was extracted, as equal to:

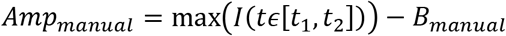

and the width at half max (WHM) was taken to be the total time that the signal was greater than the average of the:

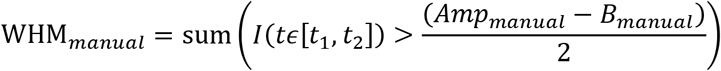

The genetic algorithm was run at default settings for 172 generations. The objective function *Err*_1_ was the sum of the squared differences of the correlation coefficients of the manual measurements and automatic measurements normalized to manual measurements. *Err*_2_ was the fraction of signals with no waves incorrectly selected to contain waves compared to the manual ground-truth annotated data:

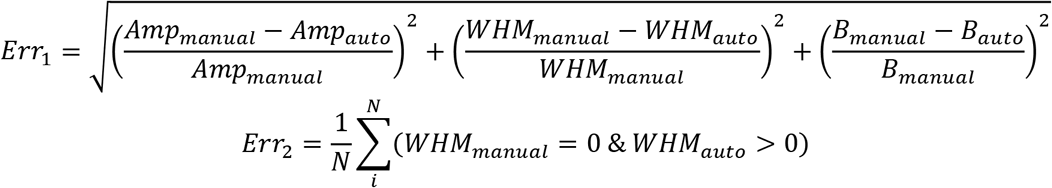

where N is the number of signals analyzed (N = 233). A Pareto front was generated to demonstrate the trade-offs between summary statistic error and false positives. Parameter values were selected that minimized both measurement error and false positives (Fig. S2, Table S2).

### In Vivo Analysis

The in vivo data was analyzed qualitatively by dividing each movie into 135 second clips and classifying each as having either no activity, spikes, ICTs, ICWs, fluttering, or being unanalyzable because of larval motion. This was done with a custom MATLAB script which divided each movie into clips. One random clip was looped at a time until all the data had been classified. Afterwards, the fraction of time spent in each regime was reported for each condition.

### Statistical Analysis

The median of ROIs in each disc for each summary statistics was compared across genetic conditions, pouch sizes, and between compartments (Fig. 2). ROIs within two spatial bins of the edge were excluded from the median to avoid edge effects. To compare across conditions, the two-tailed, unpaired, Student’s t-test was performed. Significance was verified with the Wilcoxon rank sum test, which gave similar results. To compare across compartments, medians were taken after dividing discs into A compartment and P compartment such that the boundary was a vertical line fitted to points along the A/P axis. The two-tailed, paired Student’s t-test was performed. Significance was verified with the Wilcoxon signed-rank test, which gave similar results. The F-test for model fit relative to a constant model was used to determine whether each summary statistic was related to pouch size (Fig. 3).

### Visualization

To explore the impact of spatial position, developmental progression, and genetic perturbations on Ca^2+^ signatures, we represented each disc as a spatial map, and obtained a spatial composite map for each condition. These spatial composites were mapped to a canonical representation (defined by the average pouch) based on the coordinate system transformation proposed in (Fig. S4) (Schaffter, 2014).

## Author contributions

J.J.Z., Q.W., P.A.B., and C.N. designed and conceived the study. Q.W., C.N., and D.K.S. performed ex vivo imaging. P.A.B. developed methods for feature extraction, statistical analysis, and manual classification. P.A.B. and F.H. implemented in vivo analysis pipeline. F.H. performed manual classification of in vivo data. J.C., P.L., and D.Z.C. developed automated methods for image segmentation and registration. M.L. and F.H. performed in vivo imaging. Q.W., P.A.B., and J.J.Z. analyzed data and wrote the manuscript. J.J.Z supervised the study.

## Acknowledgements

The work in this manuscript was supported in part by NIH Grant R35GM124935, NSF Awards CBET-1403887, CBET-1553826, CNS-1629914, CCF-1217906, and CCF-1617735, ND Advanced Diagnostics and Therapeutics and Harper Cancer Research Institute Research like a Champion awards (QW), Walther Cancer Foundation Interdisciplinary Interface Training Project (PB), and the Notre Dame Advanced Diagnostics & Therapeutics Berry Fellowship (CN). The authors gratefully acknowledge the Notre Dame Integrated Imaging Facility. We would like to thank Mark Alber, Christopher Paolucci, Jeffrey Kantor, Gregory Reeves, Alyssa Lesko, Jamison Jangula and Yogesh Goyal for helpful feedback, Brandon Greenawalt (Notre Dame Center for Social Research) for helpful conversations regarding statistical methodology, S. Restrepo for sharing unpublished observations, Jahmel Jordon and Kara Synder for technical support, and members of the Zartman Lab.

## Supplemental Information

Table of Contents
Table S1 Overview of lines
Table S2 Parameters of feature extraction pipeline
Table S3 SI movies
Figure S1 Signal extraction from wing discs
Figure S2 Optimization of image analysis parameters
Figure S3 Manual identification of the wing disc pouch orientation
Figure S4 Transformation of spatial maps onto a canonical axis
Figure S5 Correlation of laser power with image intensity

**Table S1.**
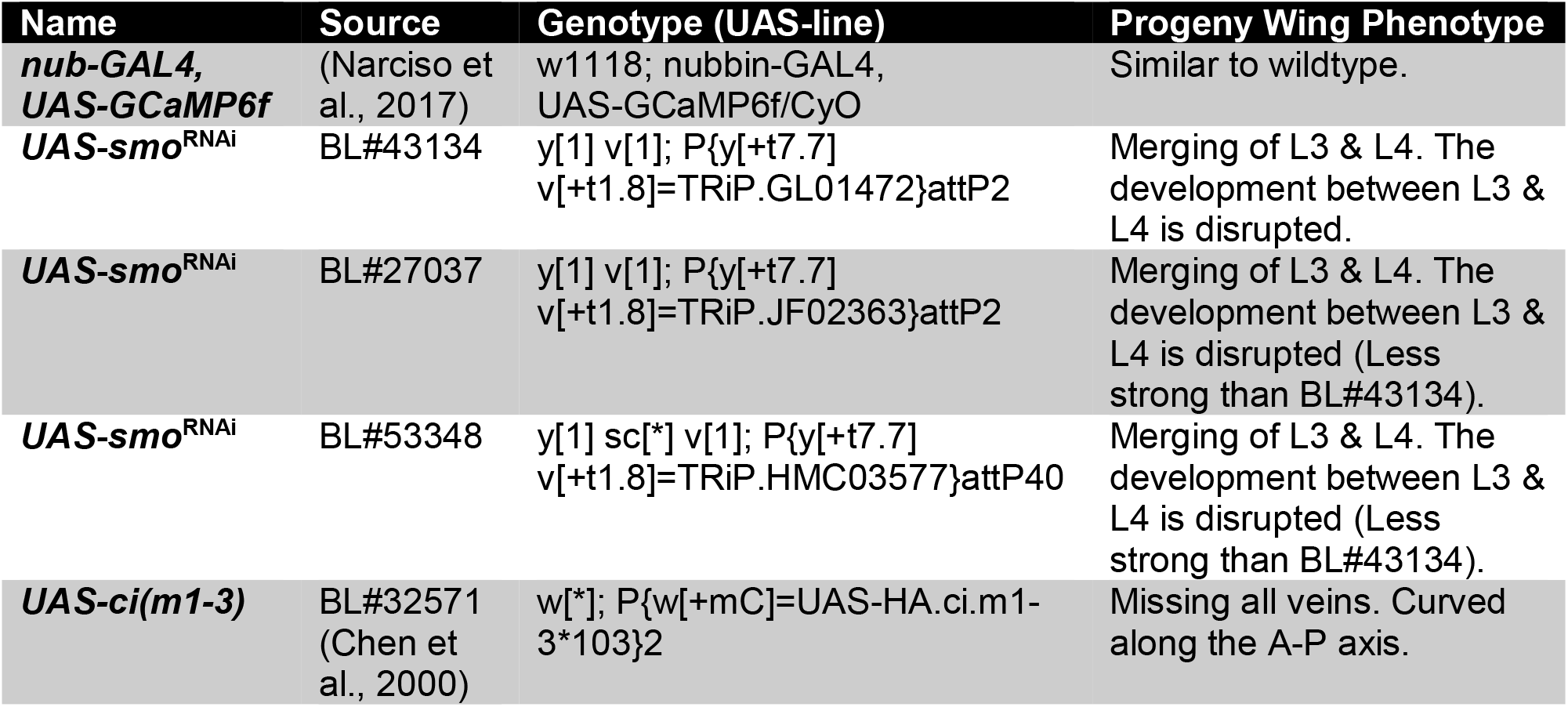
Overview of lines.

**Table S2.** Parameters of feature extraction pipeline.

| **Parameter** | **Optimized Value** | **Meaning** |
| --- | --- | --- |
| **Amp<sub>min</sub>** | 0.29 | Minimum peak amplitude |
| **WHM<sub>min</sub>** | 23 s | Minimum duration of peak |
| **Refractory Period** | 10 s | Minimum time between peaks |
| **$\sigma_{\min}$** | 8 s | Parameter for high-frequency Gaussian filter for removing noise |
| **$\sigma_{\max}$** | 1100 s | Parameter for low-frequency Gaussian filter for adapting to changes in basal level |

**Table S3.** SI movies.

| **Title** | **Description** |
| --- | --- |
| ***Movie S1*** | Intercellular calcium waves ex vivo in tester line, nub-GAL4, UAS-GCaMP6f |
| ***Movie S2*** | Intercellular calcium waves ex vivo in tester line crossed with UAS-smo<sup>RNAi</sup> |
| ***Movie S3*** | Intercellular calcium waves ex vivo in tester line crossed with UAS-ci(m1-3) |
| ***Movie S4*** | Intercellular calcium waves in vivo in tester line, nub-GAL4, UAS-GCaMP6f |
| ***Movie S5*** | Intercellular calcium waves in vivo in tester line crossed with UAS-smo<sup>RNAi</sup> |

**Figure S1.**
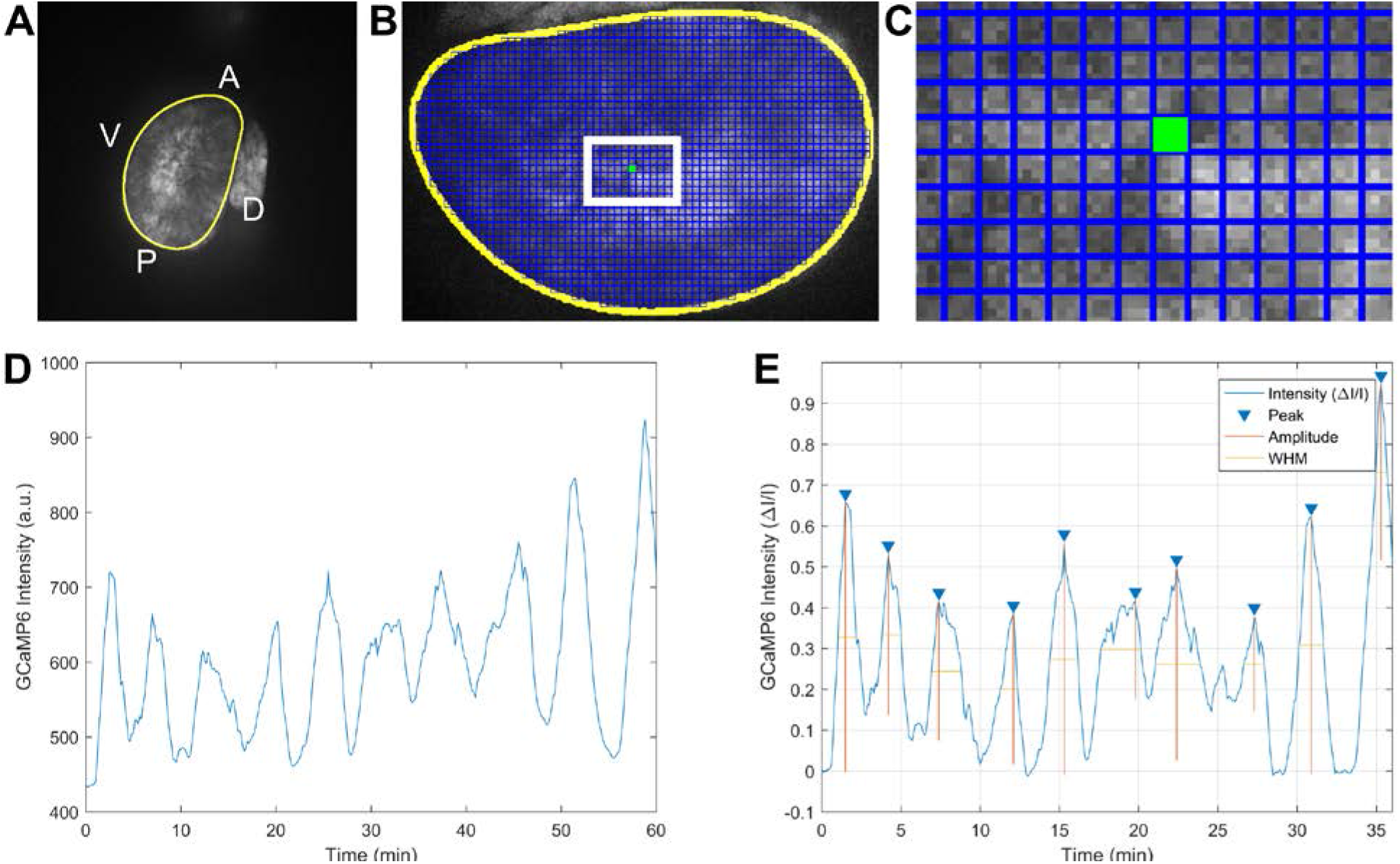
Signal extraction from wing discs. (*A*) t-slice projection of time-lapse video. (*B*) Manual mask around pouch, and grid of square regions of interest (ROIs). (*C*) Individual ROI is averaged over space to obtain a onedimensional (1D) intensity profile. (*D*) Raw intensity profile. (*E*) Normalized intensity profile with amplitudes and widths at half max (WHM) marked.

**Figure S2.**
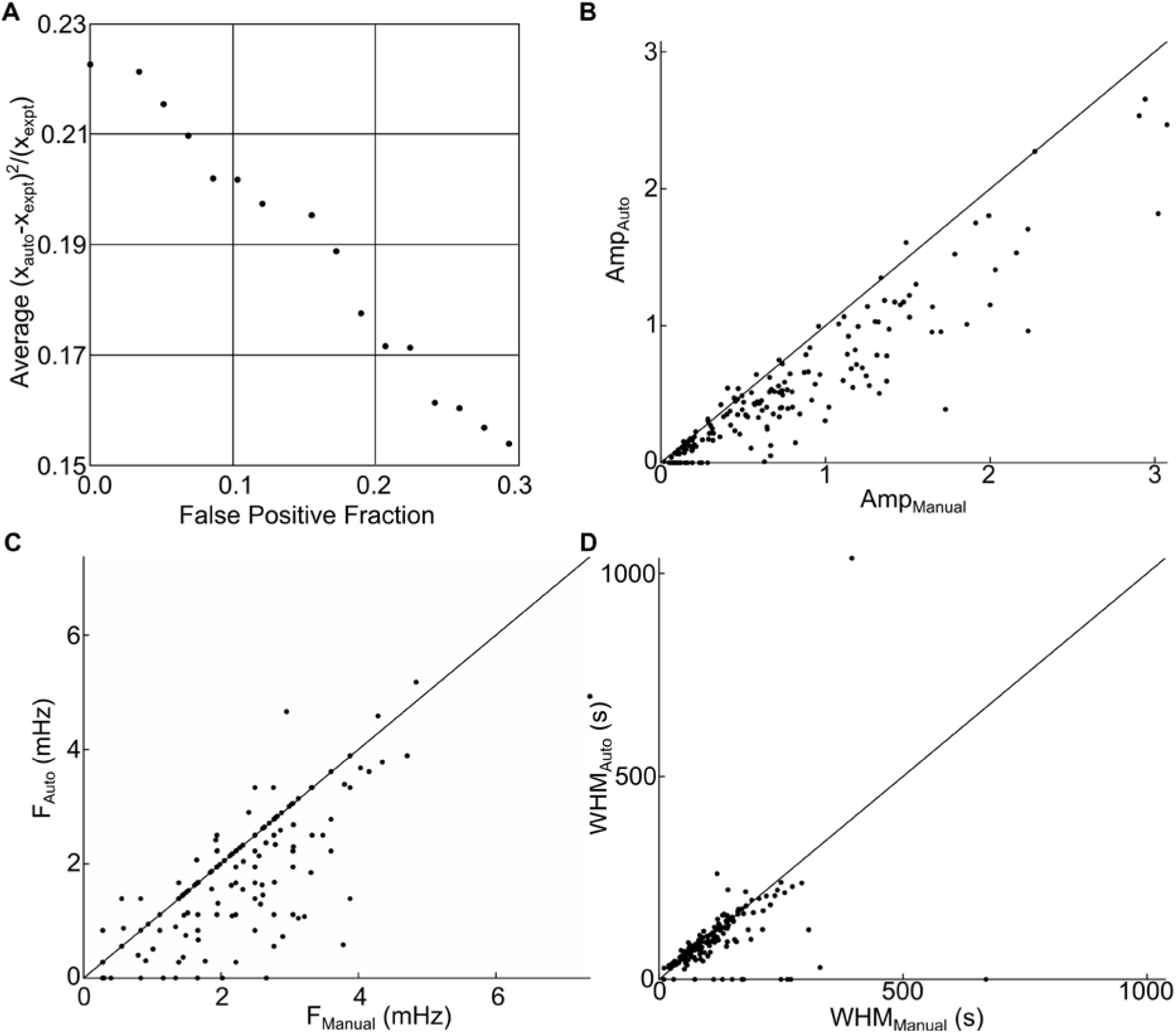
Optimization of image analysis parameters. (*A*) Pareto optimization chart for final solution. (B-D) Comparison of automatically-extracted values and manually measured values for (*B*) frequency, (*C*) amplitude, and (*D*) width at half max.

**Figure S3.**
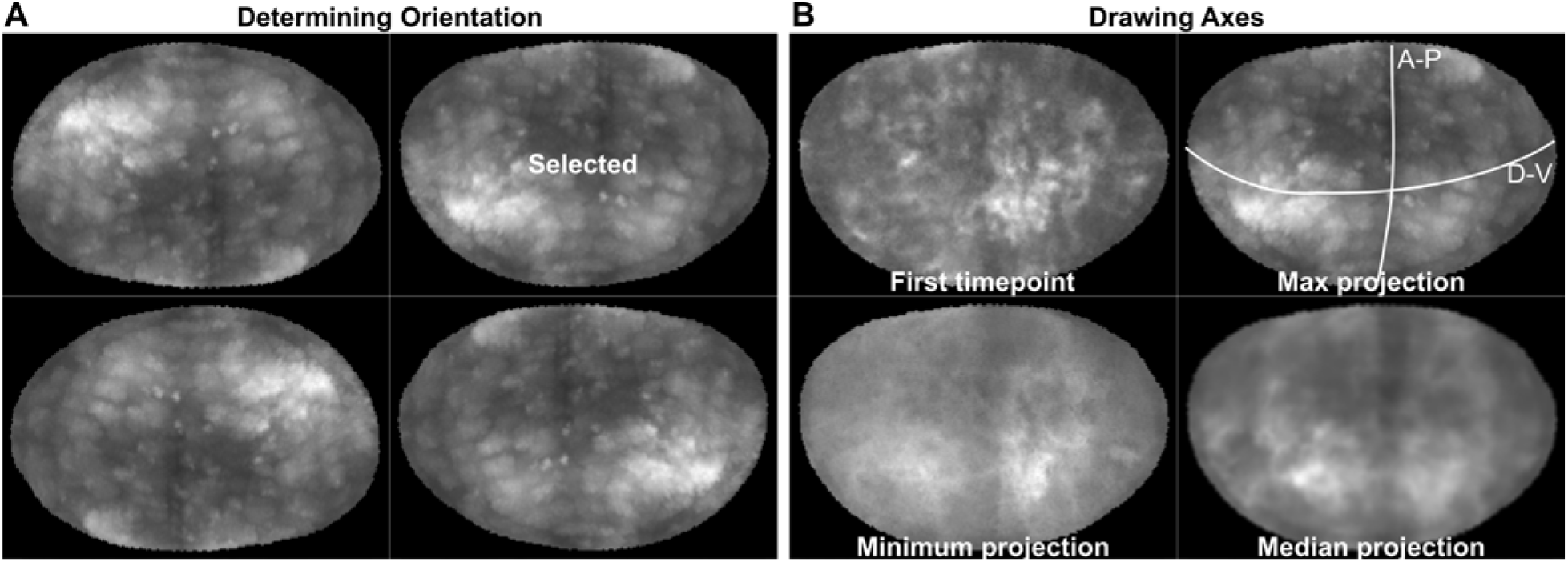
Manual identification of the wing disc pouch orientation. (A) Workflow for manual selection of orientation. Registered time stack was max projected over time in FIJI. The maximum projection was shown in a 2x2 format to the user with a non-processed, horizontally-flipped, vertically-flipped, and 180-degree rotated image presented in a random order. The user selected the image with the P compartment on the right and the D compartment on the top. Landmarks used to make the classification include the shape and size of the compartments as indicated in Fig. 1A. The pouches were presented in a random order, and were repeated at least three times. If all three initial classifications were the same, that orientation was taken to be the consensus. If more than three attempts were needed, a consensus was reached when more than half of the selected discs were the same orientation. Two discs did not reach a consensus orientation after seven attempts and were not included in the analysis. (B) Workflow for axes selection. The first timepoint, max projection, min projection, and median projection were shown to the user in a 2x2 format. Curves were drawn for the A-P and D-V axes. The A-P axis generally aligns with a sharp decrease in intensity on the maximum projection, and the D-V axis generally aligns with an increase in basal level on the minimum projection. White lines indicate manual annotations of axes.

**Figure S4.**
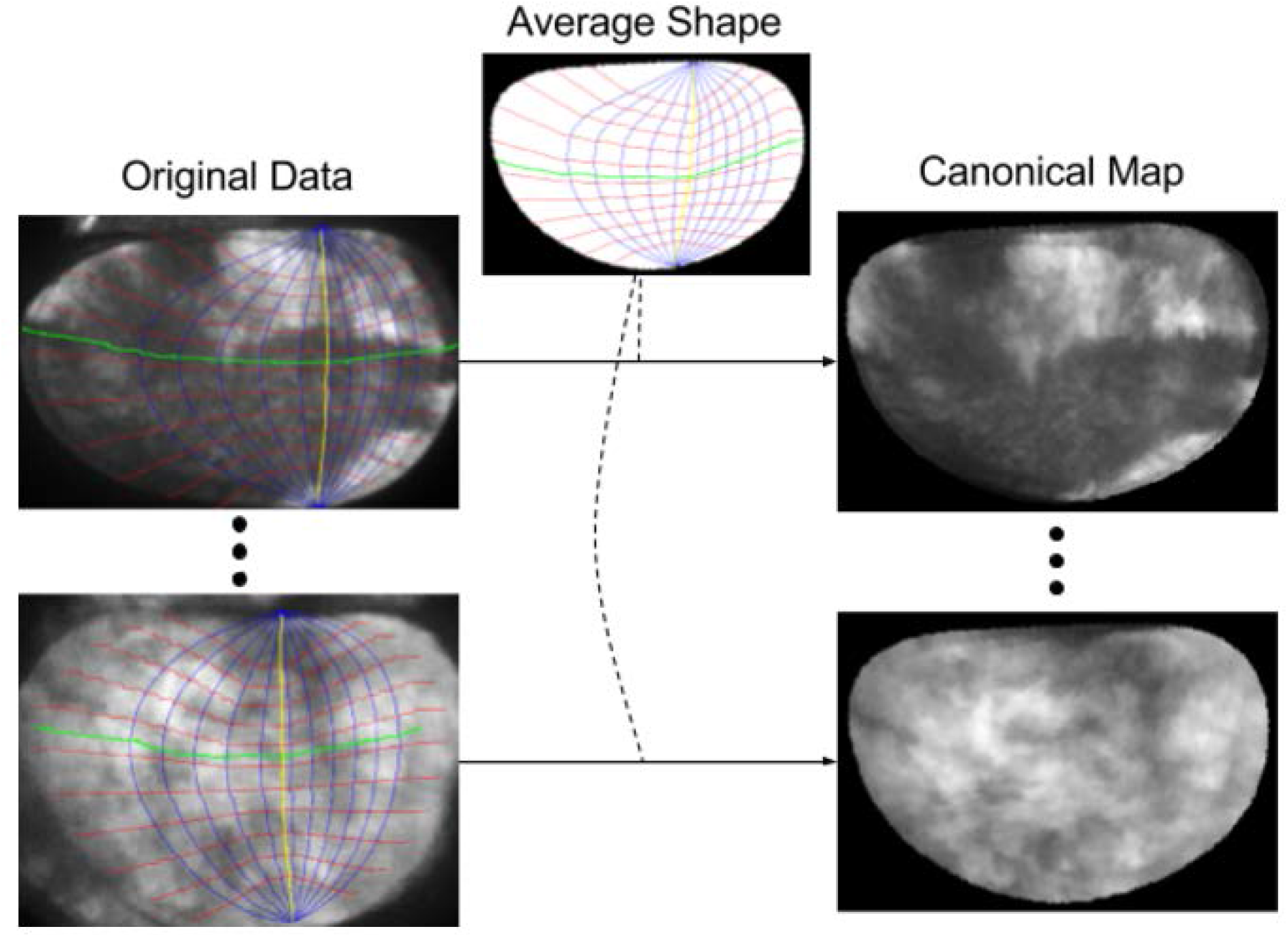
Transformation of spatial maps onto a canonical axis. A pouch coordinate system is defined along the A/P axis (the green curve) and the D/V axis (the yellow curve), similar to the latitude (cf. red curves) and longitude (cf. blue curves) of the geographic coordinate system, and is constructed for each pouch. A mapping from the pouch coordinates in an input image to the corresponding pouch coordinates of the average shape transforms the original data to a canonical map.

**Figure S5.**
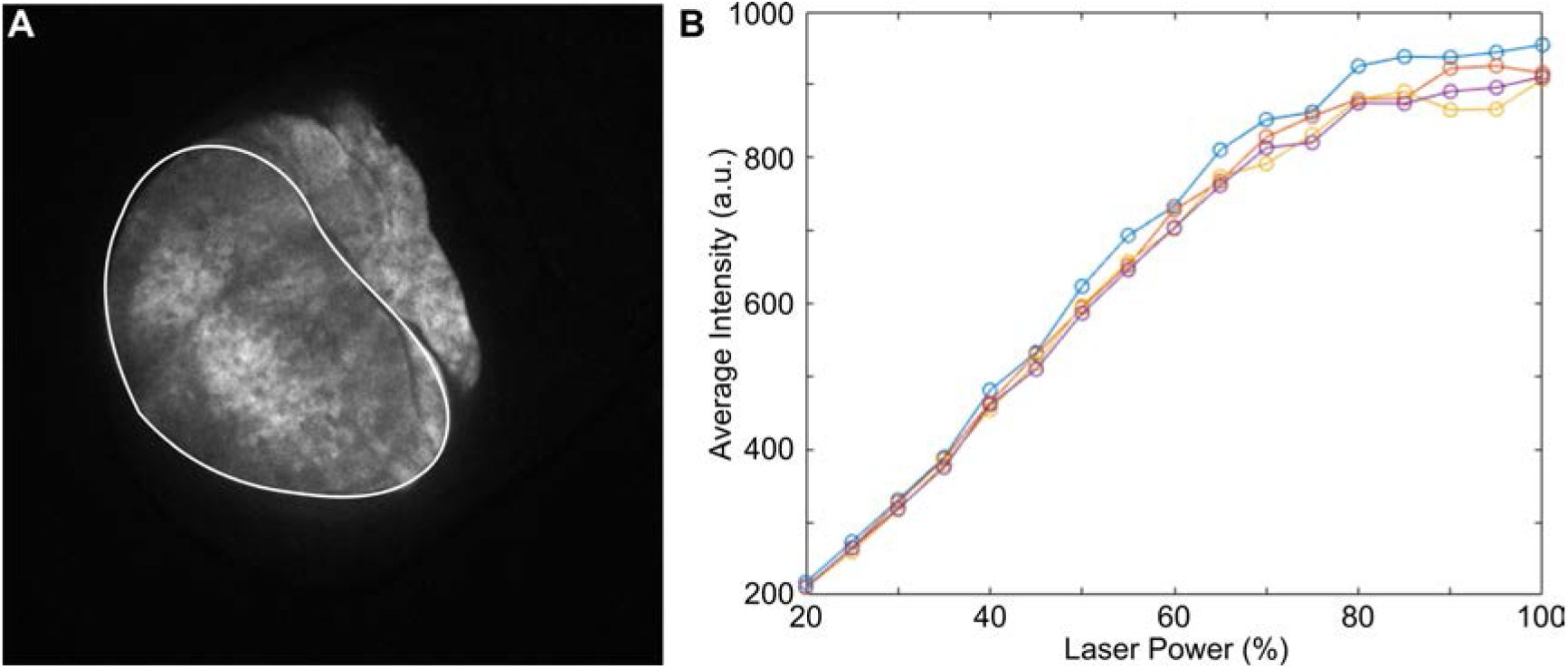
Correlation of laser power with image intensity. (A) Maximum T-projection of confocal image of nub-GCaMP6 wing disc acquired at a laser power of 50% (n-4). Pouch was segmented and the average intensity of the pouch was obtained at various laser powers. Images were acquired in a randomized order to reduce error from photobleaching. White line indicates boundary of pouch. (B) Average intensity of the pouch varies linearly with laser intensity under 80%.

